# The neutrophil enzyme myeloperoxidase directly modulates neuronal response after subarachnoid hemorrhage, a sterile injury model

**DOI:** 10.1101/627638

**Authors:** Aminata P. Coulibaly, Pinar Pezuk, Paul Varghese, William Gartman, Danielle Triebwasser, Joshua A. Kulas, Lei Liu, Mariam Syed, Petr Tvrdik, Heather Ferris, J. Javier Provencio

**Author notes:** Corresponding author: J. Javier Provencio, MD, Louise Nerancy Associate Professor in Neurology and Neuroscience, Office number: (434)297-7608.

## Abstract

Neutrophil infiltration into the central nervous system (CNS) after injury is associated with cognitive deficits. Using a murine model of aneurysmal subarachnoid hemorrhage (SAH), we elucidate the location and mode of action of neutrophils in the CNS. Following SAH, neutrophils infiltrate the meninges, and not the brain parenchyma. Mice lacking functional myeloperoxidase (MPO KO), a neutrophil enzyme, lack both the meningeal neutrophil infiltration and the cognitive deficits associated with delayed cerebral injury from SAH. The re-introduction of biologically active MPO, and its substrate hydrogen peroxide, to the cerebrospinal fluid of MPO KO mice at the time of hemorrhage restores the spatial memory deficit observed after SAH. This implicates MPO as a mediator of neuronal dysfunction in SAH. Using primary neuronal and astrocyte cultures, we demonstrate that MPO directly affects the function of both cell types. Neurons exposed to MPO and its substrate show decreased calcium activity at baseline and after stimulation with potassium chloride. In addition, MPO and its substrate lead to significant astrocyte loss in culture, a result observed in the brain after SAH as well. These results show that, in SAH, the activity of innate immune cells in the meninges modulates the activity and function of the underlying brain tissue.

## 1. Introduction

Neutrophils have been described as pathological in a multitude of neurological disorders. For example, neutrophil infiltration into the substantia nigra causes neuronal loss in mice^1^. Neutrophil adherence to cerebral capillary beds is linked to cognitive decline in a mouse model of Alzheimer’s disease^2^, and increased neutrophil infiltration in the brain after middle cerebral artery occlusion coincides with increased neuronal loss^3^. However, the cellular and molecular mechanism of action of neutrophils in the CNS is poorly understood. Our laboratory has shown that peripheral neutrophil infiltration leads to the development of cognitive deficits in a murine model of subarachnoid hemorrhage (SAH) due to loss of late long-term potentiation (L-LTP) in hippocampal neurons^4, 5^. Interestingly, although SAH neutrophils do not invade the hippocampus, they never-the-less seem to drive neuronal dysfunction^4^.

Neutrophils have a limited but potent set of effector functions that are implicated in neurological injuries. These effector functions include enzymes that catalyze the production of reactive oxygen and nitrogen species, and the degradation of extracellular matrix proteins^6, 7^. A number of these neutrophil enzymes have been linked to brain pathology after injury. For example, myeloperoxidase and elastase (both neutrophil granule enzymes) increase oxidative stress in the brain after cerebral ischemia ^8, 9^. In fact, modulation of myeloperoxidase activity after stroke leads to better behavioral outcomes^9^. Limb remote ischemic postconditioning improves stroke outcomes by decreasing the activity of NADPH oxidase and myeloperoxidase in peripheral neutrophils^10^. Inhibition of elastase improves motor recovery and spinal cord damage after injury^11^. Although all these results implicate neutrophil proteins in injury-induced neuropathology, how close the interaction needs to be between neutrophils and brain tissue, and whether these molecules directly or indirectly affect neurons, remains unclear.

Study of brain inflammatory pathology in most sterile conditions is limited due to the superimposed necrosis that causes direct brain damage and allows peripheral immune cells to invade the parenchyma thereby mixing the neuroinflammatory response and the necrosis-driven inflammatory invasion. The murine model of subarachnoid hemorrhage is an useful model to study the effect of brain inflammation on brain physiology because the direct injury occurs outside the brain in the meninges. In addition, patients with aneurysmal subarachnoid hemorrhage develop a delayed syndrome of cognitive impairment and vascular spasm^12–15^ (called cerebral vasospasm) that the murine model of subarachnoid hemorrhage recapitulates and is a readout of brain injury^16^.

In this project, we investigate the location and molecular mechanism of action of neutrophils in the central nervous system after SAH. Using a murine model of mild subarachnoid hemorrhage, we demonstrate that neutrophils affect neuronal function without invading the brain parenchyma, providing further evidence of the neuromodulatory role of the peripheral immune system in acute brain injuries. Indeed, our results also demonstrate that modulation of peripheral immune cells affects cells within the brain (i.e. astrocytes and neurons). Finally, we demonstrate that the neutrophil enzyme myeloperoxidase can directly affect neuronal function and ultimately modulates neuronal response to stimuli.

## 2. Results

### 2.1 The neutrophil enzyme myeloperoxidase is critical to the development of cognitive deficits after subarachnoid hemorrhage (SAH)

In our model of sterile brain injury, peripheral depletion of neutrophils impedes the development of spatial memory deficits after SAH^4^. To determine whether neutrophil effector function plays a role in these deficits, mice deficient in the neutrophil effector proteins myeloperoxidase, elastase and NADPH oxidase (that have been previously implicated in neuronal damage after ischemic stroke)^8, 10, 17^, were tested (Fig. 1). Both elastase and NADPH oxidase knockout (KO) mice developed delayed spatial memory deficits after SAH (Fig. 1A and 1B). On the other hand, the MPO KO mouse was protected against the development of delayed spatial memory deficits (Fig. 1C), suggesting that neutrophil derived MPO is critical to the development of this syndrome.

**Figure 1:**
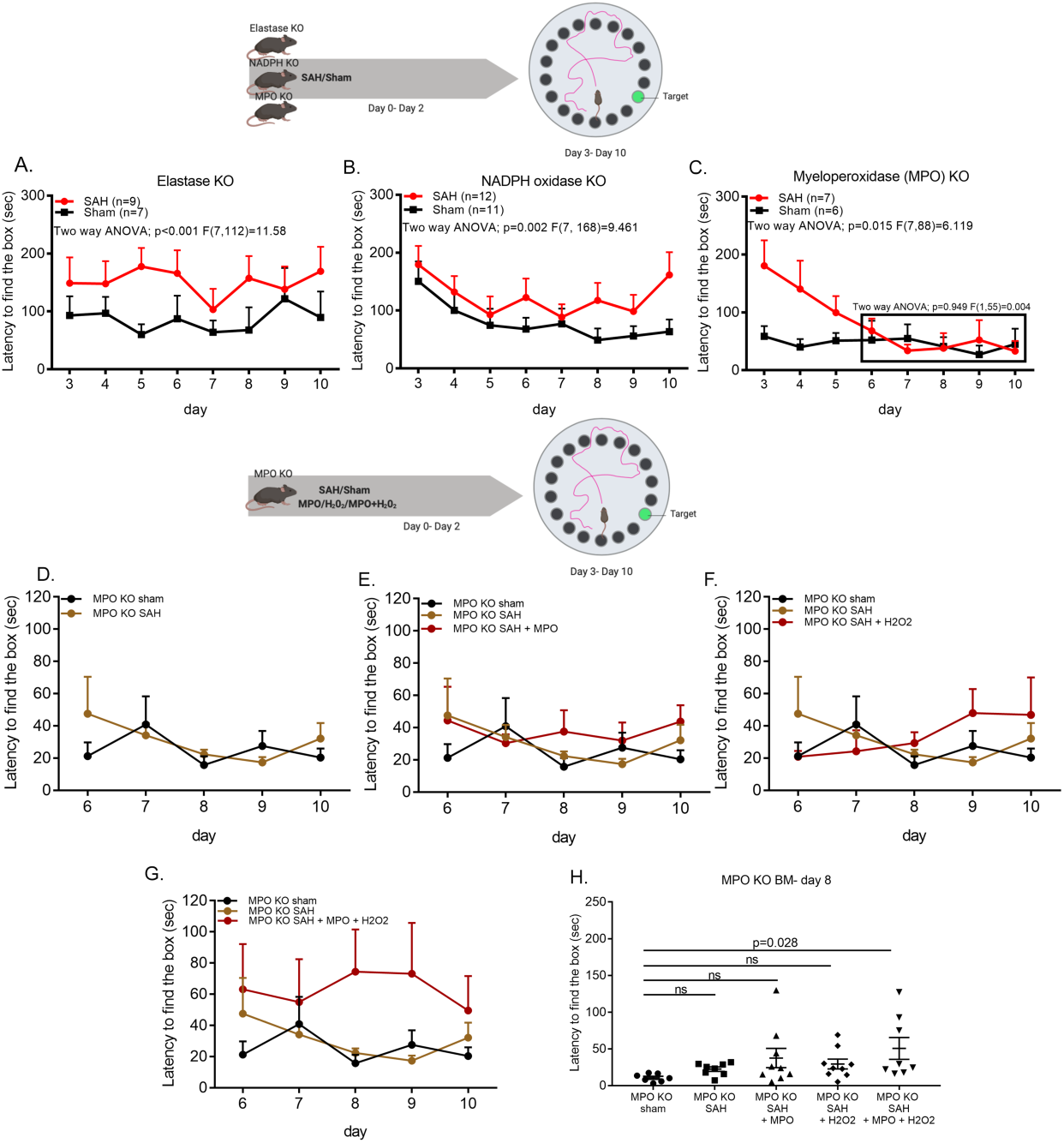
Myeloperoxidase is critical to the development of delayed cognitive deficits after SAH. Barnes maze memory task were used to determine the mechanism of action of neutrophils after SAH. Escape latency of mice lacking functional elastase, NADPH oxidase, and myeloperoxidase (MPO) were tested for 7 days starting on the 3^rd^ day after surgery in the Barnes Maze memory task (**A-C**). Mice lacking functional elastase (**A**) or NADPH oxidase (**B**) developed memory deficits after subarachnoid hemorrhage, SAH. On the other hand, MPO deficient (MPO KO) mice did not develop the delayed deficits associated with the task after SAH (**C**; boxed area). To determine whether MPO alone was responsible for the lack of deficit in the MPO KO mouse, biologically active MPO and/or H_2_O_2_ was injected into the CSF at the time of hemorrhage in the MPO KO mouse. As previously shown, MPO KO mice did not develop delayed cognitive deficits after SAH in the Barnes Maze memory task (**D**). Neither the introduction of exogenous MPO alone nor H_2_O_2_ alone in the cerebrospinal fluid (CSF) of the MPO KO mouse during SAH was sufficient to significantly affect the latency to find the goal box (**E** and **F**). The addition of MPO and H_2_O_2_ together did recapitulate the late cognitive deficit observed in the wildtype mouse (**G**). A focused analysis on day 8 after surgery showed a significance increase in latency to escape between the MPO KO sham and the MPO KO mouse with MPO and H_2_O_2_ (Student t-test: p=0.028). Diagrammatic representations of behavior experiments were created with BioRender.

To determine whether the loss of MPO activity leads to improved cognitive function, biologically active MPO was added to the CSF of MPO KO mice at the time of the hemorrhage. The addition of MPO and its substrate, hydrogen peroxide, to the MPO KO mouse recapitulates the spatial memory deficits previously present in our model (Fig. 1G and 1H). The addition of MPO alone (Fig. 1E) or hydrogen peroxide alone (Fig. 1F) was not sufficient to cause deficits suggesting that the peroxidase activity of the MPO enzyme is important for late cognitive deficits associated with SAH.

Finally, an identifying hallmark of both murine and human SAH is the presence of delayed vasospasm in large cerebral vessels^18^. In the MPO KO mouse, the diameter of the middle cerebral artery showed no spasm 6 days after SAH (supplemental figure 1).

### 2.2 Neutrophils infiltrate the meningeal parenchyma but not the brain after SAH

In order to locate the site of action of neutrophils in the CNS, the meninges and brain parenchyma were analyzed for neutrophil infiltration after SAH. Neutrophils were found in the meninges (Fig. 2) and not the brain parenchyma (supplemental fig.2, confirming our previous finding^4^) of the SAH and sham mouse. In the meninges, neutrophils are found in the venous sinuses (supplemental figure 3A) as well as the meningeal parenchyma (Fig. 2A). Although no significant changes were observed in the number of neutrophils found within the sinuses of the sham and SAH mice (supplemental figure 3B), a significantly higher number of neutrophils were observed in the meningeal parenchyma of the SAH mouse 3 days after hemorrhage compared to sham (Fig. 2B). Further characterization, using intravenous CD45 injection, showed that this increase is due to active infiltration of neutrophils into the meninges after SAH (Fig. 2C).

**Figure 2:**
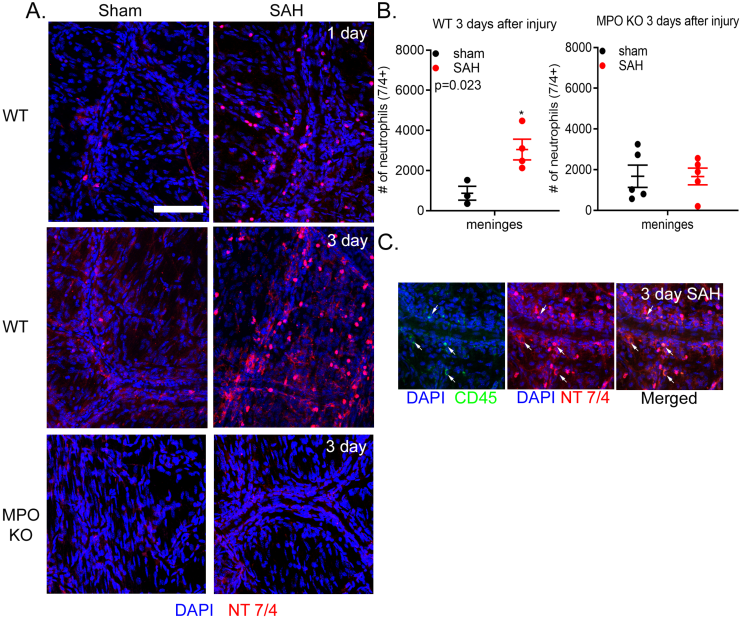
Neutrophils actively infiltrate the meningeal parenchyma after SAH. Using immunohistochemistry of whole mounted meninges, neutrophil localization within the meninges of mice was determined after SAH. Neutrophils (red; NT 7/4) were observed in the meningeal parenchyma of both sham and SAH groups of the WT and MPO KO mice (**A**). In WT mice, the number of neutrophils within the meninges increased significantly 3 days after SAH (**B**). This increase was absent in the meninges of MPO KO mice. Intravenous injection of fluorescein tagged CD45 (CD45-FITC) showed that the increased infiltration is attributed to the active recruitment of neutrophils from the periphery (**C**). Scale bar = 50 µm.

On the other hand, the MPO KO mice show few to no neutrophils in the meninges after SAH (Fig. 2A and 2B). The persistent presence of neutrophils in the meningeal venous sinuses of the MPO KO mice (Supplemental figure 3.) suggests that extravasation, rather than recruitment, may be impaired in the MPO KO. This is consistent with previous reports that show that the loss of functional MPO affects both neutrophil migration and production of key cytokines by neutrophils^19, 20^.

### 2.3 The loss of functional MPO in neutrophils skews the innate immune landscape of the CNS at baseline

The lack of functional MPO in neutrophils affects neutrophil secretion of cytokines, including IL-6, CXCL1, and MIP-1α^20^. The downregulation of these cytokines attenuates immune cell recruitment to the lungs during infection^20^. Recent evidence shows immune cells populate the meninges under normal conditions^21^. To determine whether the congenital loss of functional MPO affects the baseline immune makeup of the brain and meninges, flow cytometry was performed on both CNS compartments, brain and meninges, of the naïve MPO KO and WT mice (supplemental figure 4 gating strategy).

The MPO KO mouse has more microglia and non-microglia CD45^+^ cells in the brain than the WT mouse (supplemental figure 5A). As a proportion of non-microglia innate immune cells (CD45^+^CD11b^+^), MPO KO and WT mice had a comparable number of neutrophils in the vasculature of the brain. Analysis of monocyte populations was divided into circulating inflammatory (ly6c^hi^) and patrolling/alternative (Ly6c^lo^) subsets^22, 23^. Overall, the MPO KO mouse had more monocytes in the brain than the WT mouse (supplemental figure 5A). This increase was attributed to the high number of Ly6c^lo^ monocytes. Although the identity of these cells is unknown, we postulate that the Ly6c^lo^ population is comprised most likely of perivascular macrophages^24^.

In the meninges, a comparable number of CD45^+^ immune cells were found in the MPO KO and WT mice. No difference was observed in the number of neutrophils between the two mice, suggesting that the changes we observed in the SAH model are due to a failure to recruit peripheral neutrophils into the meninges in the MPO KO mouse. Similar to the brain, more Ly6c^lo^ monocytes populate the meninges of the MPO KO mouse at baseline.

Finally, to make sure that the baseline behavior does not differ between the WT and MPO KO mice, Barnes maze and rotarod test were performed. Although both mice have similar performance on the rotarod test, the naïve MPO KO mouse takes significantly longer to find the goal box in the Barnes maze tasks than the WT mouse contrary to the improved function seen after SAH (supplemental figure 5D). Further analyses are needed to determine if the failure to perform in the Barnes maze is due to distress (fear/anxiety) or a lack of motivation to escape.

### 2.4 MPO modulates neuronal response to stimulation

In our model, MPO appears to play a direct role in the development of the behavioral deficits associated with SAH. A possible mechanism for the development of these deficits is that MPO enters the brain to act directly on neurons after injury. Therefore, to determine whether MPO can act directly on neurons, primary neuronal cultures were generated using the Thy1GCaMP3 mice. Using the calcium signal as an indicator of activity, we characterized neuronal activity after MPO, H_2_O_2_, or MPO+H_2_O_2_ administration. The calcium traces show that the addition of MPO alone has little to no effect on neuronal activity (Fig 3A). H_2_O_2_ alone abolishes neuronal activity (Fig. 3A). Indeed, neurons exposed to H_2_O_2_ die within 2 hours (Fig. 3B). The addition of MPO with H_2_O_2_ significantly dampens neuronal activity (Fig. 3A) and results in minimal neuronal loss (Fig. 3B). This suggests that the catalysis of peroxide to reactive oxidants by MPO produces molecular species that are not immediately detrimental to neuronal survival.

**Figure 3:**
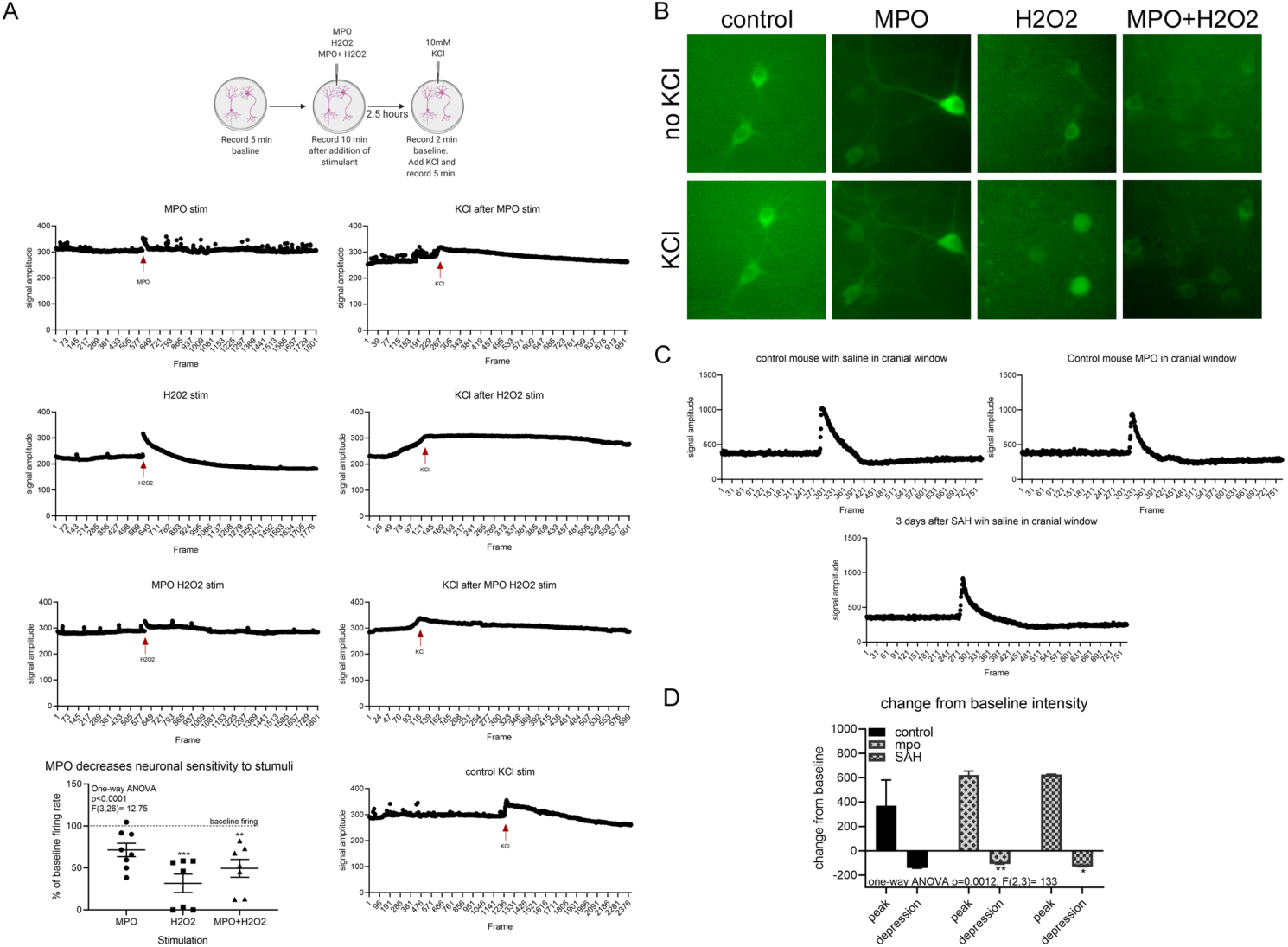
The enzymatic activity of MPO directly modulates neuronal activity. Primary neuronal cultures, obtained from Thy1-GCaMP3 mice, were incubated with MPO and/or H_2_O_2_. Time-lapsed images were taken from cultures before and after the addition of the MPO and/or H_2_O_2_. After 2.5 hours of incubation, cells were further stimulated with 10mM KCl (**A**). Representative traces of neuronal activity, as denoted by cellular calcium signal, showed the effect of MPO and/or H_2_O_2_ on baseline firing rate. MPO alone had no effect on neuronal activity, while H_2_O_2_ decreases activity by more than 75%. On the other hand, addition of MPO together with H_2_O_2_ decreased neuronal activity by 50%. Furthermore, the addition of KCl led to maximum calcium activity and subsequent recovery in all culture conditions except those treated with H_2_O_2_. Those neurons retained high level of calcium signal within the cells. The addition of MPO or MPO and H_2_O_2_ led to minimal to no neuronal death in cultures while the addition of H_2_O_2_ alone led to significant cell death (**B**). To determine whether MPO has similar effect *in vivo*, 2-photon microscopy was used on mice with SAH or exposed to MPO without SAH (**C**). Although the maximal activation of neurons was not influenced in each treatment group, SAH and the addition of MPO significantly affected the depth of signal depression observed in cortical neurons (**D**). Diagrammatic representations of experimental paradigms were created with BioRender.

Finally, to determine whether the presence of MPO affects neuronal response to stimulus, primary neurons incubated, for 2.5 hours, with MPO, H_2_O_2_, or MPO+H_2_O_2_ were stimulated with 10mM potassium chloride (KCl) solution. Neurons incubated with MPO or MPO+H_2_O_2_ show an attenuated response to KCl (Fig. 3A), further supporting the hypothesis that MPO dampens neuronal responses to stimulation.

Because the *in* vitro system does not fully mimic *in* vivo conditions, 2-photon microscopy and KCl-induced cortical spreading depolarizations (CSD) were used to determine if MPO alters neuronal response to stimuli in the live animal. Using calcium signaling as a measure of neuronal activity, we determined the peak calcium signal during CSD and the subsequent signal depression in the presence of MPO. We compared this change to the CSD induced signal and depression in the SAH mouse. Of note, because the highest level of neutrophil infiltration occurs 3 days after SAH in our model, our analysis of CSD was focused to this time point in the SAH mouse.

As demonstrated in the traces generated by calcium signaling within the neurons, under all treatment conditions, MPO or SAH, neuronal depolarization (i.e. increase in calcium signal) and depression (decrease in calcium signal) was detected within minutes of KCl administration to the cortex (Fig. 3C). Analysis of the maximum signal intensity (baseline - peak) showed no difference between groups, suggesting that MPO and SAH do not affect the depolarization potential of neurons. Analysis of depression (baseline – minimal signal) showed that both MPO administration and SAH leads to less depression in the neurons compared to controls (Fig. 3D). Because calcium signaling is more complex than a proxy for potential firing, this higher depression value may represent a failure of neurons to successfully sequester or remove all the calcium from their cytosol. It could also reflect changes in intrinsic optical properties of the affected cortical parenchyma.

To our knowledge, these experiments are the first to demonstrate that MPO has a neuromodulatory effect in sterile brain injury. However, we have been unable to locate MPO protein within the brain at any timepoint after SAH. This raises the question of contributions of intermediate cells, between the neutrophils in the meninges and the neurons in the brain parenchyma.

### 2.5 Astrocytes but not microglia are affected by meningeal derived MPO

It is unclear how neutrophils in the meninges affect neurons within the parenchyma of the brain. We investigated the possibility that MPO’s effect on neurons in the hippocampus may also act through a cell intermediary. To determine whether MPO acts on glial cells in the brain, we characterized the effect of SAH on microglia and astrocytes in the hippocampus.

Microglia morphology was characterized using Iba1 expression in the hippocampus with the understanding that infiltrating macrophages will also express Iba1 (supplemental figure 6A). Interestingly, no changes were observed in the number of Iba1^+^ cells or their fluorescent intensity in the CA1 region of the WT and MPO KO mouse at either day 3 or 6 after SAH (data not shown). To determine whether the activation phenotype of these cells was affected after SAH, the number of cell processes was quantified using a modified Scholl analysis. In the WT mouse hippocampus, no changes are observed in microglia ramification after SAH at day 3 (supplemental figure 6B). 6 days after injury, the number of processes detectable on the injured mouse decreased significantly (supplemental figure 6B). In the MPO KO mouse, at day 3 after injury, microglia lost the ramified morphology observed in the sham mouse. This change reverted to control levels 6 days after SAH (supplemental figure 6B). These results suggest the changes in the microglia population are not due to MPO but a function of the injured/inflamed state of the CNS.

Astrocytes were characterized in the CA1 region of the hippocampus by their expression of GFAP and vimentin (Fig. 4). GFAP was abundantly expressed in the CA1 sub-region of the hippocampus at day 3 (Fig. 4A) and day 6 (Fig. 4E) in both WT and MPO KO mice. At day 3, although no changes were observed in the number of GFAP^+^ astrocytes after SAH (Fig. 4C), GFAP intensity was significantly decreased in the CA1 (Fig. 4B). Furthermore, more astrocytes contained colocalized GFAP and Vimentin 3 days after SAH (Fig. 4D), suggesting increased activation of these cells^26^. At day 6, both GFAP^+^ cell number and intensity were significantly decreased after SAH (Figs. 4E-G) with no change in the colocalization of GFAP and vimentin (Fig. 4H). Interestingly, none of these changes were observed in the MPO KO mouse after SAH at either timepoint (Fig. 4A-E), suggesting astrocytes are a good candidate intermediary between the meninges and neurons in the brain.

**Figure 4:**
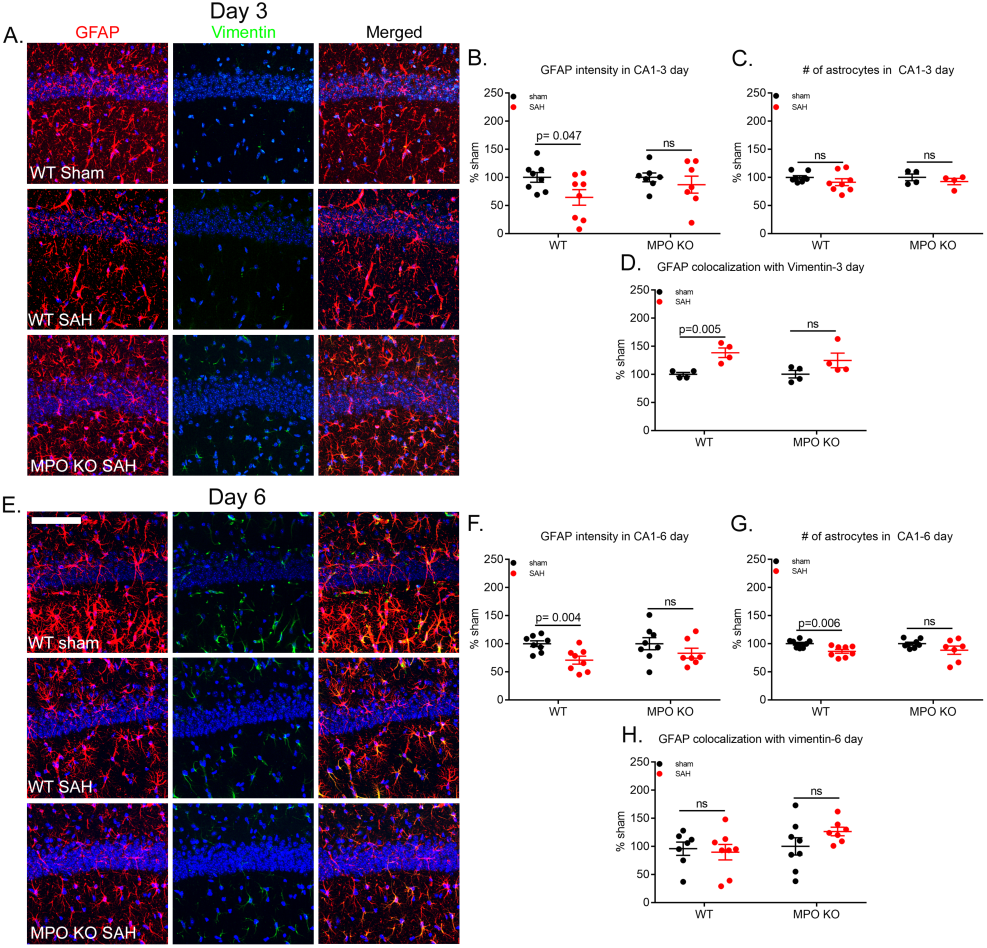
Dynamic changes in astrocytes cytoskeletal protein expression in the wildtype after SAH is absent in the MPO KO mouse. Representative confocal micrographs of GFAP (red) and Vimentin (green) positive astrocytes in the CA1 subregion of the hippocampus 3 (**A**,) and 6 days (**E**) after SAH. 3 days after surgery, GFAP intensity is significantly decreased, p=0.047, in the CA1 subregion of the WT mouse. A change that is absent in the MPO KO mouse (**B**). Further analysis showed no effect on the number of astrocytes at this time point (**C**). As expected, the injury led to a significant increase in colocalization of GFAP and vimentin in the WT mouse, p=0.005 (**C**) and not in the MPO KO mouse. 6 days after surgery, both GFAP intensity (**F**) and the number of GFAP^+^ cells (**G**) were significantly decreased in the CA1 region of the WT mouse after SAH, p=0.004 and p=0.006 respectively. No changes were observed in the colocalization of GFAP and vimentin at this time point (**H**). No changes were observed in the MPO KO mouse at either time point. Scale bar= 50µm.

### 2.6 MPO activity leads to astrocyte death in culture

The data above suggests MPO can directly affect neuron and astrocyte function within the brain. Because MPO is not detectable in the parenchyma of the brain, the proximity of astrocytes to the meninges make these cells a likely intermediary. This altered functioning of astrocytes could lead to the dysregulation of neuronal activity in the brain. To test this hypothesis, primary astrocyte cultures were stimulated with MPO, H_2_O_2_, or MPO+H_2_O_2_ for 4 hours. After which, cell morphology and survival were assessed.

Exposure to MPO alone had very little effect on astrocyte morphology and survival. However, the addition of H_2_O_2_ or MPO+H_2_O_2_ led to significant cell death in culture (supplemental figure 7), suggesting that astrocytes are especially vulnerable to the oxidative stress caused by peroxide and MPO catalytic activity. These results further support the loss of astrocytes evident in the mouse brain after SAH (Fig. 4). This may represent an interaction by which astrocytes interact with meningeal neutrophil-derived molecules at or near the glia limitans found at the subpial-brain interface.

To determine whether astrocyte derived molecules can affect neuronal function, primary neuronal culture activity was analyzed after exposure to astrocyte conditioned media. Both H_2_O_2_ and MPO with H_2_O_2_ led to astrocyte death making conditioned media unreliable as a source of secreted molecules. In addition, because necrotic death leads to the release of cytotoxic molecules, which are known to cause neuronal dysfunction, we focused on media from astrocytes cultured with MPO alone. The MPO alone conditioned media had no discernable effect on neuronal activity (data not shown), suggesting that MPO effects on the brain may be due to MPO mediated astrocyte death, leading to the dysregulation of neuronal support.

## 3. Discussion

### 3.1 Summary

The results of the present project provide clear evidence that immune activity in the meningeal compartment directly affects functions of the underlying brain. Using subarachnoid hemorrhage (SAH) as a sterile injury model, we show that through the action of its enzyme myeloperoxidase (MPO), neutrophils affect neuronal function. Our 2-photon and *in vitro* experiments demonstrate that MPO can directly affect neuronal activity, and that neuronal sensitivity to stimuli is decreased in the presence of MPO. We also find evidence that MPO modulates astrocyte survival *in vivo* and *in vitro*, although how astrocytes affect neuronal function still remains unclear. Finally, the modulation of neutrophil activity, through the removal of functional MPO, leads to significant change in the immune landscape within and around the brain. This suggests that neutrophils and myeloperoxidase likely play an important role in how immune cells populate the meninges.

The effect of MPO on behavior offers an opportunity to investigate the physiology that underlies delayed neuronal dysfunction after SAH. There are two likely routes by which MPO could mediate neuronal function: a direct effect on neurons, and/or through an intermediate cell type such as astrocytes or microglia. Histological evidence in our studies suggests against microglial activation as a mediator of MPO based neuronal dysfunction. The data in neurons and astrocytes is more promising.

### 3.2 Myeloperoxidase directly modulates neuronal activity

Previous findings from our laboratory demonstrate that SAH leads to delayed cognitive deficits in mice^5^. These deficits are attributed to the loss of long-term potentiation (LTP) in the hippocampus^4^. Changes in LTP and NMDAR subunits after SAH suggest an effect of the injury on neuronal physiology. Here, we find that MPO and MPO with H_2_O_2_ significantly decrease neuronal activity. Furthermore, our *in vivo* experiments using 2- photon microscopy, show that MPO’s activity, like SAH, decreases neuronal depression after cortical spreading depression.

These experiments show a clear modulatory effect of MPO on neuronal activity but raise an important question about how MPO gets from the meninges to hippocampal neurons. Immunohistochemistry in the brain after SAH does not show significant quantities of MPO in the brain which suggests either that there are methodological issues with MPO detection in the brain or that other mechanisms exist. These are areas of interest in our group.

In addition, the mechanism(s) through which MPO’s activity affects neurons is not clear. Because MPO has no identified, dedicated cell surface receptor, it is unclear which moieties on the neuronal membrane are being acted upon. However, we do know that MPO’s main enzymatic activity is the production of reactive oxygen species (ROS), specifically the chlorinating agent, hypochlorous acid (HOCl)^27^. ROS have been implicated in neuronal dysfunction and degeneration in a multitude of models^28^. These data most support the mechanism of MPO modulating the membrane property, i.e. receptors and channels, on the neuronal surface as evidenced by the changes in calcium imaging. Other possible mechanisms include internalization of MPO through a non-specific mechanism and subsequent intracellular activity which could lead to calcium changes from internal stores^29^.

### 3.3 Enzymatic activity of myeloperoxidase leads to impaired astrocyte function

The other possible pathway is the action of MPO on supporting brain cells that affect neuronal dysfunction. SAH in WT mice shows a significant loss of astrocytes in the hippocampus six days after SAH. This decrease is preceded by a significant decrease in GFAP coverage, which coincides with an increase in vimentin expression. These changes are absent in the MPO KO mouse, suggesting that neutrophil activity in the meninges affects astrocytes in the cortex. Astrocytes are critical to the maintenance of homeostatic balance within the nervous system. Their ability to support neurons, both structurally and through the provision of neurotransmitters, lactate, etc., is critical to the maintenance of normal neuronal function in the CNS^30^. Astrocytes also maintain the integrity of the CNS milieu by acting as a barrier, the blood-brain and blood-CSF barriers, between the circulatory system and the CNS. These barriers are especially critical during inflammation ^31, 32^. For example, the loss of GFAP positive astrocytes leads to increased leukocyte infiltration in the brain after stab injury^32^ and in EAE^31^. Although our model is devoid of CNS immune infiltration, the loss and/or impaired function of astrocytes may hinder their ability to ‘provide’ the necessary support to neurons after SAH and may represent a complimentary mechanism leading to neuronal dysfunction.

An alternate possibility for astrocyte interaction in late deficits is that astrocyte activation may represent an attempt at regeneration/recovery. Several studies have demonstrated that loss of GFAP leads to less reactivity and process elongation in astrocytes^33, 34^. Furthermore, the loss of intermediate filaments in astrocytes enables rapid regeneration after injury^34^, suggesting that the loss of GFAP after SAH may facilitate the initiation of recovery mechanisms within the CNS. Interestingly, the loss of GFAP coincides with increased hippocampal neurogenesis^35^ in the SAH mouse. SAH mice show increased neurogenesis in the subgranular zone of the hippocampus 4-7 days after SAH^36^. Therefore, the downregulation of GFAP may contribute to the generation of a proliferative/recuperative milieu in the hippocampus after SAH.

### 3.4 Peripheral immune cells modulate the immune landscape of the CNS

Recent reports have implicated peripheral and meningeal immune cells in the modulation of CNS function. Meningeal immune cells modulate output of higher order neurons as demonstrated in animal models of spatial learning, autism spectrum disorders, sickness behavior, and depression^37–40^. A recent hypothesis has designated the meninges as a tertiary lymphoid organ^41^, especially in autoimmune disease like multiple sclerosis, suggesting the meninges can become immunological under highly inflammatory conditions. However, current evidence shows the presence of resident immune cells within the meninges^21, 42^, suggesting that meninges present a local source of immune cells for the CNS after injury. In our model of SAH, there is active recruitment of neutrophils to the meninges after injury. Interestingly, the sham-treated animal meninges contain a fair number of neutrophils. The question as to whether these are resident neutrophils and the contribution of the resident population to the increase observed after injury needs to be investigated.

Neutrophils are the main source of myeloperoxidase. However, immature monocytes and microglia have been shown to express the enzyme at low levels^43^. As such, the increased number of microglia and monocytes observed in the brain and meninges of the MPO KO mouse would suggest that MPO may play a modulatory role in the activity or proliferation of these cells. Indeed, it is well established that MPO is important for the migration and extravasation of neutrophils^19, 20^. Furthermore, neutrophil derived molecules, i.e. cytokines, play a critical role in the infiltration and activity of other innate and adaptive immune cells^44^. Therefore, it is possible that the lack of neutrophils in the meninges of the MPO KO mouse significantly affect how inflamed the meninges become in response to SAH. Although neutrophils are critical for the secondary injury and deficits associated with SAH, an understanding of the role of other immune cells, like monocytes, in the central nervous system, may add to the picture of inflammation after SAH.

## 4. Conclusions

This paper provides evidence that activity of neutrophils outside the brain can modulate neuronal function during injury. We show that MPO is critical to the behavioral changes observed in SAH mice. Its effects include modulation of both astrocytic and neuronal activity within the parenchyma (Fig. 5). One possible interpretation of the data suggests that after SAH, neutrophils degranulate in the subarachnoid space, leading to activation of astrocytes that form the glia limitans in the subpial space and the blood brain barrier. Reactive oxygen species from MPO’s enzymatic activity in this space leads to the down regulation of GFAP, retraction of astrocytic processes and cellular death at this critical interface. The loss of integrity at this barrier then leads to infiltration of inflammatory mediators, including MPO into the CNS which then influences brain functions. Furthermore, MPO dampens neuronal response to stimuli, thereby dysregulating whole circuit activity. Although other mechanisms are likely at play, we propose that this dysregulation is likely a critical factor in the development of cognitive deficits associated with SAH.

**Figure 5:** Neutrophil derived myeloperoxidase directly affects brain parenchymal cells after sterile injury subarachnoid hemorrhage. The results presented in this study clearly demonstrate that myeloperoxidase, MPO, can directly modulate CNS output by either affecting neuronal firing and response to stimulation or dysregulating astrocyte function. WE postulate that 3 days after SAH, neutrophils infiltrate the meninges. Therein, they release MPO, a granule protein, into the subarachnoid space. MPO’s activity within this space leads to dysregulation and death of astrocytes at the subpial brain interface. Further leading to dysregulation of neuronal support and ultimately CNS output.

## 5. Methods

### 5.1 Animals

Young (8-12 weeks old) male mice were used in all experiments. Mice were kept on a 12 hour:12-hour light cycle at room temperature (22-25°C). Food and water were provided *ad libitum*. All experiments were done with the approval of the University of Virginia Animal Care and Use Committee.

Transgenic mice deficient in the enzymes elastase (B6.129X1-Elane tm1sds: elastase KO), myeloperoxidase (B6.129X1-Mpo tm1Lus: MPO KO) or neutrophil cytosolic factor 1(NCF- component of the NADPH oxidase complex (B6(Cg)-Bcf1 m1J: NADPH oxidase KO) (The Jackson Laboratories) were used. All transgenic mice used were on a C57BL/6 background; therefore, control experiments were conducted on C57BL/6J mice.

### 5.2 Subarachnoid hemorrhage (SAH)

SAH or sham surgeries were performed as previously reported^16^. Briefly, mice were anesthetized using isoflurane, and placed in a prone position. A 3mm incision was made on the back of the neck along the midline, the atlanto-occipital membrane was entered, and a conserved subarachnoid vein punctured. Bleeding from the vein was allowed to stop on its own. For the sham experiment, the procedure was the same except the atlanto-occipital membrane was not entered and the vein was not punctured.

### 5.3 Myeloperoxidase (MPO) injection

MPO KO mice received intracisternal injections of biologically active MPO enzyme, H_2_O_2_ or MPO with H_2_O_2_ in the setting of SAH. Biologically active MPO (ab91116; Abcam, Cambridge, MA, US) was reconstituted in sterile phosphate buffer saline (PBS) to a stock concentration of 1 µg/µl. Per product sheet recommendation, MPO production of hydroclorous acid (HOCl) requires a H_2_O_2_ concentration of 0.0012% per unit of activity. Mice were given a cisterna magna injection of either 0.8 µg of MPO, 0.8 µg MPO with 0.0012% H_2_O_2_ in PBS, or 0.0012% H_2_0_2_ in PBS at the time of surgery.

### 5.4 CD45-FITC injection

Mice were injected with 3 µg of anti-CD45-FITC antibody (Thermofisher, Waltham, MA, US) intravenously 30 minutes prior to euthanasia. Mice were then perfused with PBS and 4% paraformaldehyde. Meninges were dissected, stained for neutrophils with the anti-neutrophil antibody 7/4-FITC (Abcam, Cambridge, MA, US; 1:100), and imaged using an Olympus FV1200 confocal microscope and Fluoview software.

### 5.5 Flow cytometry

Flow cytometry was performed on both the brain parenchyma and meninges of WT and MPO KO naïve mice. To generate a single cell suspension for flow cytometry, the meninges and brains were collected and processed as previously described^45^. Briefly, meninges and brain samples were dissected and processed in DMEM media with 10% bovine serum albumin (BSA). Samples were digested using 1 mg/ml DNAse and 1.4 U/ml collagenase in Hanks Balanced Salt Solution with magnesium and calcium. Samples were then incubated in a 37°C water bath, triturated and strained through a 70 µm cell filter. Meningeal samples were then centrifuged and resuspended in DMEM media with 10% BSA and kept on ice until staining. Brain samples were further processed using a 40% Percoll solution in PBS to remove the myelin debris. Following Percoll, samples were rinsed with DMEM with 10% BSA and kept on ice until staining. Once single cell suspensions were obtained, samples were blocked with Fc block and incubated with an antibody cocktail containing Ly6G-FITC, Ly6C-PeCy7, CD11b-efluor 780 (Life Technologies, Carlsbad, CA, US; 1:200), CD45-Pacific blue (Biolegend, San Diego, CA, US; 1:200), and fixable viability dye-506 (Life Technologies, Carlsbad, CA, US; 1:1000) for 30 minutes at 4°C. Samples were then analyzed using a Gallios cytometer (Beckman Coulter, Brea, CA, US), and FlowJo software (FlowJo, LLC, Ashland, OR, US).

### 5.6 Immunohistochemistry

Mice were transcardially perfused with 4% paraformaldehyde in PBS. Brains and meninges were dissected and processed separately for immunohistochemistry. Brains were post-fixed in 4% paraformaldehyde for 24 hours followed by cryoprotection in a 30% sucrose solution. Brains were then frozen and 30 µm sections collected using a cryo-microtome. Sections containing the hippocampus were selected, rinsed in PBS, and blocked in a 0.3% Triton-X with 5% normal goat serum solution. Sections were then incubated with rabbit anti-Iba1 (Wako Chemicals, Japan 1:500), anti-neutrophil antibody 7/4-FITC (anti-Ly6B) (Abcam Cambridge, MA, US; 1:100), chicken anti-GFAP (Abcam, Cambridge, MA, US; 1:1000), and rabbit anti-vimentin (Abcam, Cambridge, MA, US; 1:200) overnight at room temperature. The following day, sections were rinsed and incubated with AlexaFluor conjugated secondary antibody (ThermoFisher Scientific, Waltham, MA, US; 1:500) for 1 hour at room temperature, mounted on Superfrost Plus slides (ThermoFisher, Waltham, MA, US), and cover-slipped using Vectashield mounting medium with DAPI (Vector laboratories, Burlingame, CA, US).

Meninges were removed from the skullcap and kept in PBS until processing. Non-specific binding sites were blocked using PBS containing 1% BSA, 5% normal goat serum, 0.1% Triton-X, 0.2% tween, and Fc block^46^. Meninges were then incubated in a primary antibody cocktail containing the anti-neutrophil antibody 7/4-FITC (Abcam, Cambridge, MA, US; 1:100) at 4°C overnight.

### 5.7 Confocal image acquisition and analysis

All images were obtained on the Olympus FV1200 confocal microscope with Fluoview software. Each image was collected as a Z-stack. Prior to image analysis, all images were converted to a maximum intensity image by collapsing all stacks using ImageJ/Fiji (NIH, Bethesda, MD, US) and blinded for analysis. Image Analysis was then performed by a technician/student in the laboratory not aware of the treatment conditions.

Brain: Images of both the left and right CA1 regions of the hippocampus were obtained using a 20x oil immersion objective. Composite images containing the nuclear marker DAPI and the cellular stain of interest (Iba1 for microglia and GFAP/vimentin for astrocytes) were then generated. For cell counts, the multipoint tool was used to count all cell within an image. For fluorescent intensity, confocal images were converted to 8- bit images. Then using the measure tool, the fluorescence intensity was determined for each image. Counts for both left and right hippocampi were added and divided by 2 to generate an average, number of cells or fluorescence intensity, per case. Finally, using the ‘Coloc 2’ application for ImageJ/Fiji, a co-localization index for GFAP and Vimentin was obtained for each hippocampus. The values were also averaged for left and right hippocampi to generate a value for each case.

Meninges: In order to image both the meninges containing the transverse sinus and the meningeal parenchyma (fibrous portion), whole mounts of the meninges were stained for neutrophils (NT 7/4; Abcam, Cambridge, MA, US). Using 20x oil immersion objective, a 3 × 3 tile was obtained of the transverse sinus and the meninges. Each image was obtained as a Z-stack. Prior to analysis, a maximum intensity image was obtained for each stack, and a DAPI and NT 7/4 composite generated. To randomize data acquisition, a 9 × 9 grid was placed on each image within the tile and the number of neutrophils counted within every 6^th^ square (starting from the left top square, 6 squares were counted (5 horizontally and 1 vertically)). As such, a total of 7 squares were counted per grid laid images for a total of 252 um^2^ area (36 um^2^ area/grid square) counted out of 4225 um^2^ area (image size). The number of cells was then added across all 9 images (from 3 × 3 tile) per case, divided by 252 (to generate # of cells/ um^2^), and multiplied by 4225 to generate a representative number of neutrophils present in each case.

### 5.8 Microglia Scholl analysis

Brain images of sections labelled with anti-iba1 antibody, were converted to 8-bit files. The iba1 fluorescence threshold of the images was set at 90% intensity, and a 10 × 10 grid was placed on each image. To randomize data acquisition and minimize bias, the fifth row of squares was cropped out and analyzed. Cells were only included in the analysis if fully located within the selected row. The ‘Scholl analysis’ application was then used to quantify the number of processes observed on selected cells using the protocol previously described ^47^.

### 5.9 Vasospasm

Previous studies from our lab show the presence of vascular spasm in the middle cerebral artery (MCA) of SAH mice early (at day 1) and, more importantly, in a delayed manner (6 days after the hemorrhage) corresponding to DCI in mice^5^. Since the peak neutrophil infiltration occurs in our model 3 days post-hemorrhage and delayed vasospasm is correlated with DCI, we focused our vasospasm analysis on the delayed vasospasm occurring on the 6^th^ day after SAH.

The arterial tree of each mouse was labeled using the vascular dye Microfil (Flow Tech, Carver, MA), and the cross-sectional MCA diameter was measured. Briefly, animals were anesthetized using sodium pentobarbital, transcardially perfused with 20 ml of cold PBS followed by 20 ml of cold 1% paraformaldehyde in PBS, then injected with 10 ml of Microfil dye. Brains were dissected, rinsed in PBS, then cleared using methyl salicylate (Sigma-Aldrich, St Louis MO US). The ventral surface of each brain was then imaged using a Leica DM 2000 LED light microscope. Because vasospasm leads to non-uniform constriction of arteries, there are affected and unaffected areas. The percent constriction was calculated as a ratio of the smallest diameter to the largest cross-sectional diameter of the MCA within a 2 mm segment distal to the posterior wall of the internal carotid using ImageJ and converted to a percentage.

### 5.10 Behavior analysis

Prior to surgery, mice were randomly assigned to treatment group using a random number generator. In addition, treatment conditions were blinded to the person administering the behavioral test.

#### Barnes Maze

Previous studies show that mice with SAH develop deficits in performance on the Barnes maze task 6-8 days after the hemorrhage^5^. Therefore, in the present study, this task was used to assess the presence of cognitive deficits after SAH. Briefly, mice were habituated to the maze for 2 days before the surgery. Habituation was performed by placing mice in the middle of the maze and leaving them to explore for 5 min. Testing started on day three post-surgery and continued for 7 consecutive days thereafter. During testing, mice were placed in the middle of the maze and given a total of 300 seconds to find the escape hole. Each trial ended when the mouse entered the escape hole, stayed in the entry zone of the escape hole (defined by a circle of 5 cm diameter around the escape hole) for 5 consecutive seconds, or the 300 second maximum had elapsed. All trials were recorded and analyzed using the tracking software Ethovision 13 (Noldus, Leesburg, VA, US).

#### Rotarod

To determine whether the naïve MPO KO and WT mice showed different motor coordination and learning, mice motor performance was analyzed using the rotarod test. During this test, only the latency to fall was analyzed. In brief, mice were placed on the rod at low speed and left to acclimate for 60 seconds. Mice were then tested with a continuous increased speed on the rod for 5 consecutive minutes. Experiments ended when mice fell of the rod or the 5 min have elapsed. Each mouse was given 3 unique trials separated by a 30 minutes rest period in the home cage.

### 5.11 Primary cultures

#### Primary cortical neurons

Neuronal cultures were established using *Thy1-GCaMP3* neonates as previous described^48, 49^. Briefly, cortices of P0 neonates were dissociated, in minimal essential medium (MEM) with 10% heat inactivated FBS and penicillin/streptomycin, using a 1000ul micropipette. Cells were filtered using a 70um cell filter and seeded on sterile 12 well plates coated with poly-D-lysine (0.1 mg/ml). Two hours after seeding, media was changed to neurobasal A media with B27, glutamax and penicillin/streptomycin. 25% of cultures media was subsequently exchanged every two days. Seven days in vitro (DIV) cultures were incubated with either 0.8 ug of MPO with or without H_2_O_2_ (0.0012%) or H_2_O_2_ alone. All cultures conditioned were imaged using a widefield Leica DM6000B microscope outfitted with a CCD camera. Before stimulation, a 5-minute time-lapse baseline video (at a rate of 2 frames per second) was obtained for each culture.

Following the 5 min baseline recoding, neurons were stimulated with either MPO with or without H_2_O_2_ or H_2_O_2_ alone, followed by a 10minutes time-lapse recording. Cultures were left to acclimate to the stimulus for 2.5 hours. Cells were then stimulated with 10mM potassium chloride (KCl) solution. The first 10 min of the KCl stimulation were also recorded. Prior to analysis, all time-lapse recordings were blinded. Using Imaris (Bitplane, Concord, USA), 8-10 cells were chosen at random and a maximum fluorescent intensity value was generated for each cell at each frame acquired in the time lapse video. The values were plotted on excel and firing rate (peaks) were counted before and after stimulation in each experimental group (control, MPO, MPO+H2O2, and H2O2). Percent baseline values were then determined by dividing the firing rate of each cell after the stimulation to the firing rate before the stimulation. Values were averaged across each cell to generate a single value for that time lapse recording.

#### Primary astrocyte culture

Primary astrocyte cultures were established as previously described^50^. In brief, brain cortex was collected from P0 pups, dissociated in 2.5% Trypsin in HBSS solution, and vigorously pipetted using a 10ml serological pipette to generate a single cell suspension. Cells were then spun and resuspended in astrocyte media (DMEM, high glucose, 10% heat inactivated FBS, and penicillin/streptomycin). Cells were then platted on T75 culture flask and incubated at 37 degrees. 2 day after platting, media was changed and every 3 days thereafter. On the 7^th^ day DIV, culture flask was shaken and rinsed to remove microglia and oligodendrocyte progenitor cells. The generated pure astrocyte culture was further incubated for 12-14 days, after which the culture was split. 2 days after the split, cultures were detached, rinsed and transferred to 15mm culture dish for testing. As with the primary neuronal culture, astrocytes cultures were imaged, exposed to 3 experimental conditions, MPO, MPO with H_2_O_2_, or H_2_O_2_ alone, then incubated for 4 hours. At the end of the incubation period, cultures were imaged. Cells were counted in the before and after images to determine the toxic/morphological effect of each experimental condition on astrocytes.

Of note, for both neuron and astrocyte cultures, each mouse brain was plated on 12 culture dishes (or 1 12-well culture plate). Triplicates of each experimental condition (control, MPO, MPO+H2O2, and H2O2) were performed for each mouse. For analysis, data was collected from each triplicate, and averaged to generate a single data point for a given mouse brain.

### 5.12 2-photon imaging of neuronal calcium activity

All imaging experiments were performed on a Olympus FVMPE-RS system equipped with sensitive GaAsP detectors and a resonant scanner. A cranial window was prepared on Thy1-GCaMP3 mice as previously described^51^. In brief, mice were anaesthetized, mounted on the stereotactic instrument, and the skin on the scalp removed to expose the skull. A 3mm diameter circle was made in the parietal bone with a microdrill 2 mm lateral and 2 mm caudal to the bregma. The edge of the circle was further thinned by drilling until the circled bone cap was able to be flipped and removed with fine-tipped forceps. A few drops of 0.9% NaCl saline solution was applied on top of the exposed dura mater. A small area of dura was carefully cut with a fine needle and a solution of drug (MPO or saline) applied directly to the cortex. The cranial window was covered with a glass coverslip, sealed at the edges with super glue. To facilitate the injection of KCl into the cortex, a 1mm diameter burr hole was prepared 3 mm caudal and lateral to the cranial window. A flattened nail (used as a handle to orient mouse head during imaging) was horizontally glued the contralateral occipital bone.

The cranial window implanted mice were kept under anesthesia for the duration of the imaging session. 3 unique regions were imaged within the cranial window separated by a 10 minutes interval. Images were acquired using a Olympus 25X NA 1.05 objective lens with an 920 nm infrared laser. The line averaging on the resonant scanner was set to 6 to obtain images with 512X512 resolution and 2.5 Hz acquisition speed. A target field of view was selected at a depth between 100 nm and 200 nm and continuous 5 minutes-long T-series imaging session was recorded.

To induce KCl-triggered spreading depolarization, a 20 um diameter pulled micropipette was filled with 1 M KCl, mounted on a nanoliter injector (WPI) and inserted into the burr hole. For imaging, a 1-minute baseline was first recorded. Followed by cortical spreading depression generated by the application of 100 nL of 1 M KCl to the cortex though the burr hole, and 4 additional minutes of recording.

Each generated file was blinded and then transferred to the Imaris software (Bitplane, Concord, USA). Within each image, 10 individual neurons were selected at random. Maximum fluorescent intensity was collected for each cell at every frame during the recording. Values were averaged across all cells and the 3 unique field of view for each mouse to generate a single data point for each given mouse. Finally, increase fluorescence from baseline (peak and depression) were calculated across groups to determine whether the addition of the MPO, saline or SAH affected the neuronal response in cortical spreading depression.

### 5.13 Statistics

Graphpad Prism 8.1 (Graphpad, La Jolla, CA, US) software was used to analyze all of the data obtained. Student’s t-test or analysis of variance (ANOVA) were used to determine whether differences between treatment groups were statistically significant. Significance was attributed to p values less than 0.05.

## Author contributions

JJP conceived of the study and supervised the design and performance of the experiments. APC contributed to the research design and performed the majority of the experiments and data analysis presented. APC also wrote the manuscript under the guidance and supervision of JJP. PP contributed the original research design of the concept presented in this paper, performed and analyzed the experiments using the transgenic mice presented in the paper. PV performed all MPO injected behavioral experiments presented in the paper. WG performed the microglia Scholl analysis experiments, including staining and data analyses. DT conducted the behavioral experiments on the naïve mice. JAK and HF established the primary astrocyte cultures and were instrumental in the design of the astrocyte cultured experiment designs. LL and PT performed the 2-photon experiments presented. MS read and edited the manuscript.

## Acknowledgments

This work was supported by NIH (NINDS) 1RO1NS074997 awarded to JJP. Special thanks are extended to Dr. J. Kipnis lab for his advice on experimental design, and the Kipnis lab for help while conducting this project, specifically to Drs A. Louveau and S. Gadani for their help with the meninges and flow cytometry experiments, respectively. Thanks to Drs. John R. Lukens and Kevin S. Lee for review of the manuscript.

---

**Supplemental figure 1: MPO KO mice lack vasospasm in the middle cerebral artery after subarachnoid hemorrhage.** Left panel denotes the region of middle cerebral artery analyzed for vasospasm (pink arrow). Confirming previous results (Altay et al. 2009), wildtype mice showed a significant decrease in vessel diameter 6 days after SAH (Student t-test: p=0.038). No significant change in vessel diameter was observed in MPO KO mice after SAH.

**Supplemental figure 2: Neutrophils do not infiltrate the brain after SAH.** Immunohistochemical analysis of neutrophils (green; Ly6B) in the hippocampus of the SAH mouse at 3 days, showed no neutrophils in the brain parenchyma and parenchymal blood vessels (red; CD31). Scale bar = 250µm.

**Supplemental figure 3: Neutrophils populate the dural sinuses of both WT and MPO KO mouse after SAH.** Representative micrographs of neutrophil (red; NT 7/4) localization within the transverse sinus of the dura at days 1 and 3 after injury in the WT and MPO KO mouse (**A**). No significant increase was observed in the number of neutrophils in the sinus of WT and MPO KO mouse after SAH (**B**). Scale bar =50µm.

**Supplemental figure 4: Flow cytometry gating strategy for brain and meningeal tissue preparation.** Data collected was first gated on live cells, followed by gates set on singlets and single cells. In the brain, microglia cells were identified as CD45^int^ and CD11B^hi^. Using the ‘not gate’ function on Flow Jo, the remaining non-microglia CD45^+^ cells were then analyzed for monocytes and neutrophil proportions (**A**). In the meninges, CD11B^+^ cells were sub-gated from the CD45^+^ cell population. Of those, CD45^+^CD11B^+^, monocytes and neutrophils proportions were quantified (**B**).

**Supplemental figure 5: Loss of functional MPO leads to significant change in the innate immune landscape of the CNS.** To determine the baseline immune profile of WT and MPO KO mouse, flow cytometry analysis of the brain and meninges of the naïve mouse was performed. At baseline, more microglia and non-microglia immune cells are found within the brain of the MPO KO mouse (**A**). The increase in the number of non-microglia cells can be attributed to the increase levels of Ly6C^lo^ monocytes within the brain of the MPO KO mouse. No difference was present in the number of neutrophils in this compartment. Looking at the meninges, overall no changes were observed in the number of CD45^+^ cells in this compartment. Similar to the brain, More Ly6C^lo^ monocytes populate the meninges of the naïve MPO KO mouse. Because the functional output of this study was done using the spatial memory task, it was critical to determine the baseline behavioral performance of these transgenic mice. Interestingly, the MPO KO naïve mouse took longer to find the goal box in the Barnes Maze memory task contrary to the improved function seen after SAH (**B**). The WT and MPO KO naïve mouse show no difference in motor performance on the rotarod (**C**).

**Supplemental figure 6: Iba1^+^ myeloid cells (microglia and tissue macrophages) in the CA1 region of the hippocampus do not appear activated after SAH in either WT or MPO KO mice.** Representative confocal micrographs depicting the morphology and density of Iba1^+^ cells (green) in sham and SAH mice 3 and 6 day post-surgery(**A**). Scale bar = 50 µm. Scholl analysis of the activation status of Iba1^+^ cells (cells have fewer processes when more activated) of both sham and SAH mice (**B**). Three days after surgery, similar number of processes was observed on Iba1^+^ cells of the B6 SAH (n=4) and sham (n=4) mice (p=0.119). By day 6, there was more ramification in the SAH (n=4) mice suggesting the Iba1^+^ cells are less activated than sham (n=4) mice, p=0.004. In the MPO KO mice, at day 3, SAH (n=4) led to fewer ramifications but the variability in the sham (n=4) group precludes statistical significance, p=0.058. No changes were observed in the number of processes observed in MPO KO mouse 6 days after SAH (n=4) when compared to the MPO KO sham (n=4) group, p=0.897.

**Supplemental figure 7: Enzymatic activity of MPO leads to dysregulation and cell death of astrocytes.** To determine the effect of MPO and its substrate H_2_O_2_ astrocyte cultures were incubated with MPO and/or H_2_O_2_ for 4 hours. Surface coverage (suggesting healthy confluence of cell growth) and cell death were then assessed. Left panel represents brightfield micrographs of astrocyte cultures under the different conditions tested. As observed, the addition of MPO alone had very little to no effect on astrocytes in culture. The addition of H_2_O_2_ led to astrocytic death within the cultures. In addition, H_2_O_2_ led to complete astrocyte loss in culture. The addition of MPO and H_2_O_2_ led to changes in the surface coverage by astrocytes and the loss of 25% of cultured cells. Diagrammatic representation of experiment was created with BioRender.

