## Supplementary figures and images for "The neutrophil enzyme myeloperoxidase directly modulates neuronal response after subarachnoid hemorrhage, a sterile injury model"

### Supplemental Figure7

#### Supplemental Figure 7

A.

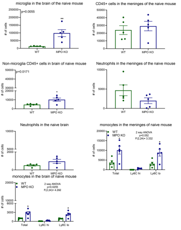


B.

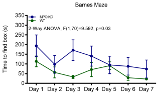


C.

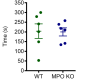

### Supplemental Figure 1

#### Supplemental Figure 1

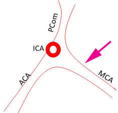


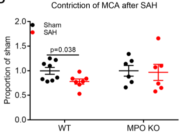

### Supplemental Figure 2

#### Supplemental Figure 2

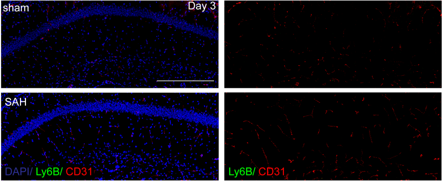

### Supplemental Figure 3

#### Supplemental Figure 3

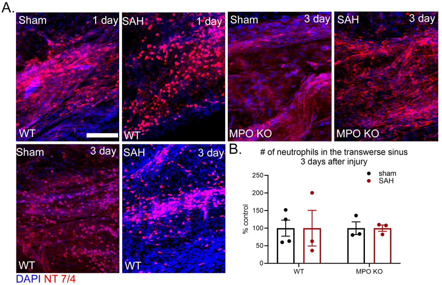

### Supplemental Figure 4

Supplemental Figure 4

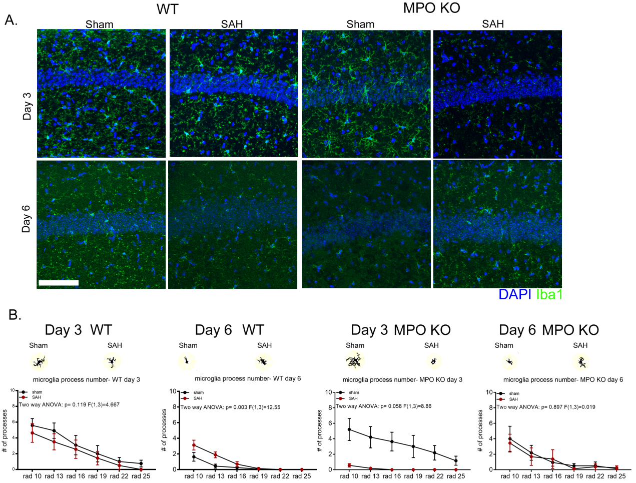

### Supplemental Figure 5

Supplemental Figure 5

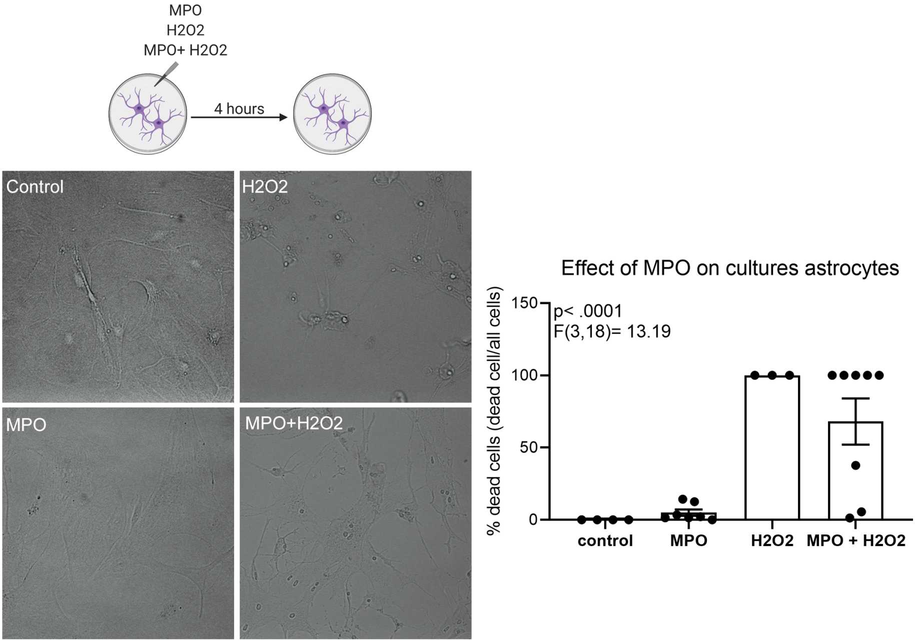

### Supplemental Figure 6

Supplemental Figure 6

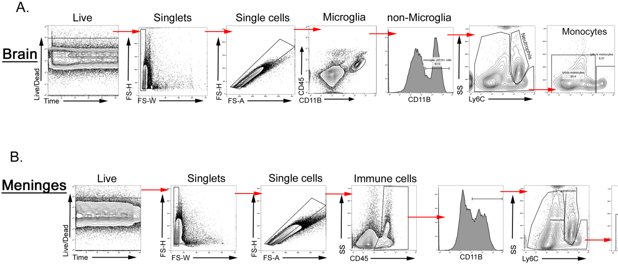
